# Sarcoptic mange; an emerging threat to Chilean wild mammals?

**DOI:** 10.1101/2020.03.02.974188

**Authors:** Diego Montecino-Latorre, Constanza Napolitano, Cristóbal Briceño, Marcela M. Uhart

## Abstract

Sarcoptic mange has been anecdotally reported in Chilean wildlife during the last decade. Although this disease can have devastating outcomes for biodiversity, there is no comprehensive assessment of this potential threat in Chile. Because the current capacity to monitor sarcoptic mange and other wildlife diseases is limited in this country, we used abnormal alopecia to search for suspect cases across several sources of information to identify, to the extent available data allow, the location and temporal trends of wild mammals with this characteristic across Chile. We surveyed park rangers, examined rehabilitation center databases, and collated citizen and media reports. The information gathered suggests that observations of alopecic wild mammals, mainly foxes (*Lycalopex* sp.), their presence in the country, and the number of species fulfilling our case definition have increased over the last 15 years. Moreover, abnormally alopecic mammals are currently located broadly in Chile. We also confirmed the utility of abnormal alopecia to define a suspect sarcoptic mange case in the most commonly affected group, *Lycalopex* foxes. Our findings are highly concerning from a conservation perspective. We thus emphasize the need for an urgent surveillance and management plan for sarcoptic mange and other diseases that may be threatening Chilean biodiversity.

## Introduction

Human-mediated global environmental change and movement of pathogens and domestic hosts have driven the emergence of disease in wildlife across the globe (Daszak et al., 2000; Tompkins et al., 2015). Evidence supporting that emerging diseases can lead to population declines and extinctions of wild species, and to subsequent changes at the community and ecosystem levels has accumulated in the last decades (Harvell et al., 2019; O’Hanlon et al., 2018; Schultz et al., 2016). Indeed, emerging disease events have become a concern for the conservation of marine and terrestrial biodiversity globally (Daszak et al., 2000; Harvell et al., 2019, 1999; O’Hanlon et al., 2018; Schultz et al., 2016; Tompkins et al., 2015).

Although disease emergence in wild species of South America is underrepresented in the scientific literature (Tompkins et al., 2015), this continent is not exempt from the global trend. For example, a cetacean morbillivirus has recently been associated with mass mortality in Guiana dolphins (*Sotalia guianensis*) in Brazil (Groch et al., 2018) and *Corynebacterium pseudotuberculosis* is currently affecting the endangered huemul deer (*Hippocamelus bisulcus*) in Chile (Morales et al., 2017).

Further, the globally emerging sarcoptic mange (Astorga et al., 2018), caused by the burrowing mite *Sarcoptes scabiei*, has been anecdotally reported in wild mammals of Chile (Alvarado et al., 2004; Verdugo et al., 2016), including within protected areas (PAs; Cunazza and Diaz, 2014; López Jiménez, 2018). Worryingly, this skin disease can have catastrophic effects on wild populations as seen in other places around the world (Martin et al., 2017). It is therefore of concern that Chilean stakeholders increasingly perceive alopecic native mammals, a classic presentation of *S*. *scabiei* infestation, as more

common and more broadly spread across the country. However, with no formal national system in place to monitor wildlife health in Chile and no centralized repository of information on disease, tracking sarcoptic mange (or other diseases) over relevant temporal and spatial scales becomes a very difficult task. Moreover, efforts to generate such data country-wide have not been made. These deficiencies and patchy data inherently limit the capability to confirm perceptions and effectively inform policy. If sarcoptic mange is a growing problem in Chile, then public and private action to prevent biodiversity impacts is warranted.

Here, we aimed to describe the trend of sarcoptic mange in Chilean wildlife over a 15-year period (2004-2018) within PAs as well as the location of cases in non-protected landscapes at the country level in recent times (2012-2019). To fulfill these objectives, we relied on abnormal hair loss (herein “abnormal alopecia”), the highly visible main characteristic of sarcoptic mange (Bornstein et al., 2001; Niedringhaus et al., 2019; Pence and Ueckermann, 2002), to establish a definition of a suspect case. With this case definition, we tracked potentially *S*. *scabiei* infested mammals over time in PAs of Chile. In addition, we combined multiple independent sources of information to also identify abnormally alopecic wild mammals outside PAs and summarize their locations in this country. Finally, we assessed the validity of abnormal alopecia to track sarcoptic mange cases by testing individuals with this condition admitted to rehabilitation centers. We discuss the implications of our findings for mammal conservation and the need for a National Wildlife Health Surveillance Program in Chile.

## Material and methods

Procedures involving human participants were limited to anonymous surveys conducted in accordance with the ethical standards of the University of California, Davis, Institutional Review Board (Protocol # 1320102-1).

### a) Description of abnormal alopecia in wild mammals within protected areas between 2004-2018

Through an anonymous survey (S1) we collected information on wild mammals observed with abnormal alopecia within Chilean public PAs between 2004-2018. Selected PAs included the 78 National Parks, National Reserves, and National Monuments with permanent park-ranger presence (by the time of the survey there were 105 PAs in total, nationwide) which represent ∼60% of the surface covered with PAs in the country. The web-based survey was sent through the Chilean National Forest Service (known as “CONAF” from the Spanish “Corporación Nacional Forestal”) to the head-ranger of each PA. Specifically, the survey inquired about live and dead wild mammals sighted or found with alopecia (yes-no) between 2004-2008, 2009-2013, and 2014-2018 in the PA. We also requested the number of individuals per species sighted with this condition per time period within the categories: 1-5, 5-20, and more than 20 individuals. The survey included pictures of Chilean wild mammals with abnormal, sarcoptic mange-like hair loss to avoid reports of animals undergoing seasonal molt (S1). Prior to deployment, the survey was improved with voluntary feedback from 2 rangers. The survey was available for response over the last 2 months of 2018. If the surveyed head-rangers had not been in that position for the 15-year period covered by this study, we requested them to submit the historical information of the corresponding PA based on internal records.

### b) Description of abnormal alopecia in wild mammals outside of Chilean protected areas

To construct a summary of the location of abnormally alopecic Chilean wild mammals outside of the PAs in recent times, we collected cases reported by citizens; cases present in digital news outlets, social media, and others (hereafter “internet cases”); and cases admitted to wildlife rehabilitation centers. To obtain information from citizens, we built a web-based platform that allowed to anonymously report abnormally alopecic wild mammals across the country (www.salud-silvestre.uchile.cl). We communicated the existence of this website and its purpose to appropriate stakeholders, and we created a social media webpage (“Salud Silvestre Chile”) to create awareness of this project. Examples of abnormal alopecic mammals were provided in the awareness campaign (S2). Reports required details on the species sighted, the date and place (District and Province) of the sighting, and at least one picture of the affected animal. This picture allowed confirmation of the species and actual abnormal alopecia by the authors. Reports not including at least one picture, or with pictures not showing alopecia, or with alopecia not resembling abnormal hair loss (e.g., seasonal molting) were disregarded. Here we included reports received between January 2017 through August 2019.

We collected data from the internet by searching for suspect (abnormal alopecia) or confirmed cases of sarcoptic mange in wildlife. We used the Google search engine with combinations of the keywords “zorro”, “guanaco”, “vicuña”, “enfermedad”, “sarna”, and “Chile”; and also “fox”, “guanaco”, “vicuna”, “disease”, “mange”, and “Chile”, the corresponding English words. We specifically targeted members of the genus *Lycalopex* (Neotropic foxes) and South American camelids (guanaco and vicuna; *Lama guanicoe* and *Vicugna vicugna*, respectively) because these were the most commonly reported wild species with abnormal alopecia according to the data from the PA survey and wildlife rehabilitation centers (see below). We included all the records with data newer than December 31st, 2003 up to November 30th, 2019.

Finally, we used a previously compiled dataset from five wildlife rehabilitation centers in Chile that covered the period 2011-2015 (Romero et al., 2019) and requested new information from these centers and an additional one to include updated data. Specifically, we solicited records from mammals admitted since 2016 fulfilling the case definition of abnormal alopecia. For each case, we asked for the geographic origin, species affected, and the date of capture or admittance. We also asked whether the etiology of the skin disease was confirmed and solicited relevant clinical information, such as the extent and severity of skin/hair lesions, body condition, and the presence of ectoparasites (e.g., fleas and ticks) at the initial examination.

For these three last data sources, we included only abnormally alopecic wild mammals that were not observed or found within the PAs that participated in the survey to avoid duplication.

### c) Validity of abnormal alopecia to track cases of sarcoptic mange

To assess the validity of abnormal alopecia to track sarcoptic mange cases, we requested skin scrapes from animals with abnormal alopecia admitted to the rehabilitation centers mentioned above from April 2016 until February 2019. Samples retrieved were assessed for the presence of mites. *S*. *scabiei* was confirmed microscopically in mite-positive samples following standard guidelines (Fain, 1968). Further, one mite was selected from each positive animal for DNA extraction and genetic confirmation. Genomic DNA was extracted using a NucleoSpin Tissue kit (Macherel-Nagel) following the manufacturer’s instructions. We conducted PCR to amplify the ITS-2 gene (450 bp [base pairs] fragment) using primers RIB-18 and RIB-3 and conditions described in Zahler et al. (1999) and the 16S rRNA gene (360 bp fragment) using primers 16SD1 and 16SD2 and conditions described in Walton et al. (2004). The PCR products were electrophoresed in 1.5% agarose gel stained with Sypro Red Protein Gel Stain. PCR fragments were sequenced in both directions at Catholic University Sequencing Unit (Santiago, Chile), inspected and aligned using software Geneious 11.0.5 (Kearse et al., 2012). Sequences were compared to those available in GenBank (National Center for Biotechnology Information, National Institutes of Health, Bethesda, Maryland, USA) using the Basic Local Alignment Search Tool to confirm the species.

Once the morphological and/or molecular identification of the mite allowed a definitive diagnosis of sarcoptic mange in the wild mammals that fulfilled our case definition (and assuming equivalent detection probability), we assessed the likelihood of *S*. *scabiei* infestation in these individuals under different levels of “true prevalence” of this mite. For this purpose, we assumed a binomial distribution parameterized by *n*, the number of animals fulfilling our case definition that were admitted to the rehabilitation centers and PCR-tested for sarcoptic mange, and 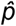, the “true prevalence” (in proportion) of *S*. *scabiei* infestation in wild mammals with abnormal alopecia. We used R (R Core Team, 2015) to obtain the probability to get x number of individuals positive to sarcoptic mange out of the *n* individuals fulfilling the case definition given a 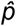 value.

The datasets are available at https://figshare.com/s/7c99a6f010278f8325e9.

## Results

### a) Description of abnormal alopecia in wild mammals within protected areas between 2004-2018

We received responses from 53 of the 78 PAs with a permanent ranger (67.95%). We excluded Juan Fernandez Archipelago National Park because it is not in continental Chile. Therefore, we report the results from the remaining 52 PAs, which cover ∼20% of the total Chilean protected surface, and whose location is shown in S3. Examples of alopecic wild mammals observed by the rangers are shown in S4. A total of 13 PAs (25.80%) across all Zones in Chile reported the presence of wild mammals, dead (n=6, 11.53%) or alive (n=13, 25.80%), with abnormal alopecia at least once between 2004-2018 (for a visualization of the Zones see S5). Specifically, these observations occurred in 3, 2, 4, and 4 PAs located in the North, North-Central, Central, and Austral Zones, respectively. Four of the 48 PAs that existed in 2004-2008 were positive (8.33%). Three of these areas are located in Central Chile while the remaining one is located in the Austral Zone. Six of the 52 PAs existing during 2009-2013 were positive (11.76%). These PAs were located in the North (n=2), North-Central (n=1), and Central Zones (n=3). Four of these PAs were new positives (0.078 incidence proportion). Finally, abnormally alopecic wild mammals, dead or alive, were observed in 11 of the 52 PAs existing during 2014-2018 (21.15%) ranging from the North to the Austral Zones of the country, except in the South Zone (3 in the North, 2 in the North-Central, 2 in the Central, 0 in the South, and 4 in the Austral Zones). Six of them were new positives with respect to the previous period (0.115 incidence proportion). If we only consider the 48 PAs that were created prior to 2004, then percentages of affected PAs were: 8.33% (n=4), 12.5%, (n=6) and 20.83% (n=10) per consecutive period of time, whilst the incidence proportion for the last 2 periods was 0.083 and 0.104, respectively. Six of the 13 PAs with cases reported more than one species affected (2004-2018). The location of PAs with reports of abnormally alopecic wild mammals, dead or alive, per time period are shown in Fig 1A. To facilitate visualization, this figure highlights the Provinces containing PAs cases, but we acknowledge that cases were not necessarily present across the entire provincial territories. Moreover, a summary with the number of PAs with reported cases per period of time and per species is shown in Table 1.

**Table 1.** Number of protected areas where wild mammals were reported with abnormal alopecia per Zone and period of time.

| Species (common name) | Zone | Number of protected areas with abnormally alopecic individuals |  |  |
| --- | --- | --- | --- | --- |
|  |  | 2004-2008<sup>1</sup> | 2009-2013<sup>2</sup> | 2014-2018<sup>3</sup> |
| *Lycalopex culpaeus* (Andean fox) | North | 0 | 1<sup>b</sup> | 0 |
|  | North-Central | 0 | 0 | 1<sup>d</sup> |
|  | Central | 2<sup>a</sup> | 3 | 2 |
|  | Austral | 0 | 0 | 2<sup>f,h</sup> |
| *Lycalopex griseus* (Chilla fox) | North-Central | 0 | 0 | 1<sup>d</sup> |
|  | Central | 2<sup>a</sup> | 0 | 0 |
|  | Austral | 0 | 0 | 1<sup>g</sup> |
| *Lama guanicoe* (Guanaco) | North | 0 | 1<sup>b</sup> | 1<sup>e</sup> |
|  | North-Central | 0 | 1<sup>c</sup> | 0 |
|  | Austral | 0 | 0 | 1<sup>f</sup> |
| *Vicugna vicugna* (Vicuna) | North | 0 | 1 | 3<sup>e</sup> |
|  | North-Central | 0 | 1<sup>c</sup> | 1 |
| *Pudu puda* (Pudu) | Austral | 1 | 0 | 1 |
| *Thylamys elegans* (Yaca) | Central | 1<sup>a</sup> | 0 | 0 |
| *Conepatus chinga* (Molina's hog-nosed skunk) | Austral | 0 | 0 | 1<sup>g</sup> |
| *Hippocamelus bisulcus* (Huemul deer) | Austral | 0 | 0 | 1<sup>h</sup> |
| **Total number of PAs** |  | 4 | 6 | 11 |

Common letter superscripts show protected areas with cases of individuals belonging to more than one species.
<sup>1</sup>: The total number of protected areas for this period was 48.
<sup>2</sup>: The total number of protected areas for this period was 52.
<sup>3</sup>: The total number of protected areas for this period was 52.

**Fig 1.**
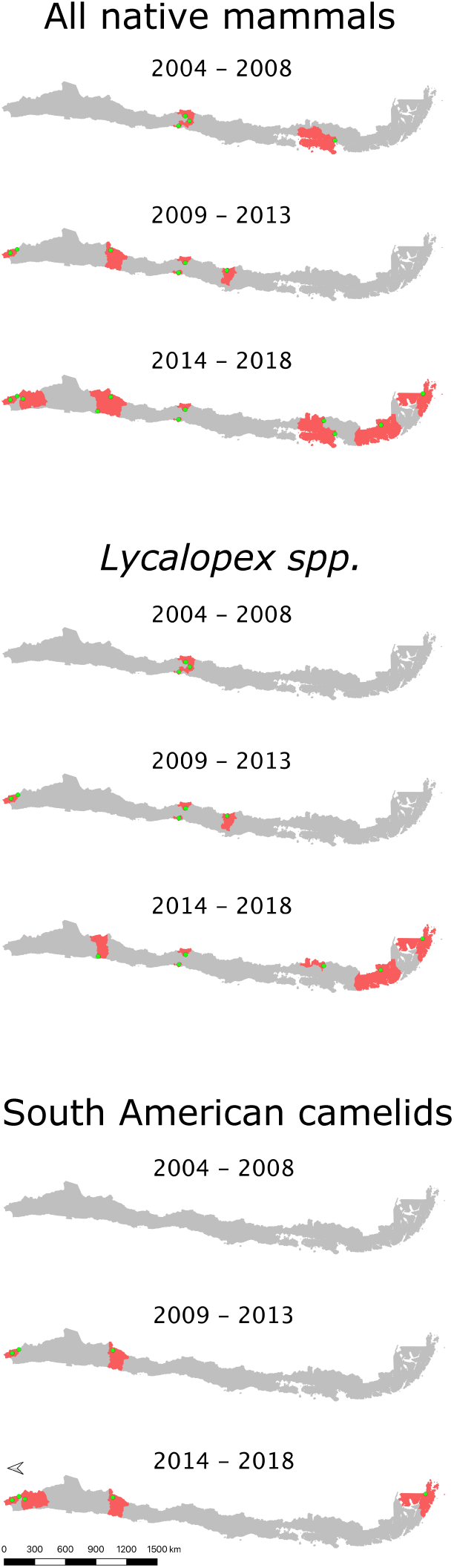
A-C. Location of abnormally alopecic native mammals, *Lycalopex* foxes, and South American camelids respectively, within protected areas (green dots) during three time periods. The red polygons show the Provinces containing protected areas with records of alopecic wild mammals over time. We present the information at this scale to facilitate visualization.

The most frequently reported native mammal species were foxes (*Lycalopex sp*.) and South American camelids (guanaco; *Lama guanicoe* and vicuna; *Vicugna vicugna*). Alopecic Chilla foxes (*L*. *griseous*) and Andean foxes (*L*. *culpaeus*) were observed in 9 of the 52 PA (17.31%; 4 and 7 PAs per species, respectively). These PAs are located across Zones except in the South (4 and 7 PAs per species, respectively) while *L*. *guanicoe* and *V*. *vicugna* were reported in 5 PAs in the North, North-Central and Austral Zones (3 and 4 total PAs per species, respectively) out of the total 52 (9.62%). Provinces containing PAs with reports for both of these taxa per time period are shown in Fig. 1B-C. The reports of these two groups (foxes and South American camelids) across 2004-2018 follow the general pattern: the number of PAs with abnormally alopecic individuals increased over time (Fig 1A). Park rangers also reported an abnormally alopecic huemul in the Austral Zone, which is probably an individual affected by *C*. *pseudotuberculosis* (Morales et al., 2017). The abundance categories reported were 1-5 affected animals in most cases, though 5–20 affected South American camelids were reported twice during the two most recent time periods (2009-2013 and 2014-2018; Table 2). A summary of the number of reported cases for all mammal species per period of time and per Zone is shown in Table 2.

**Table 2.**
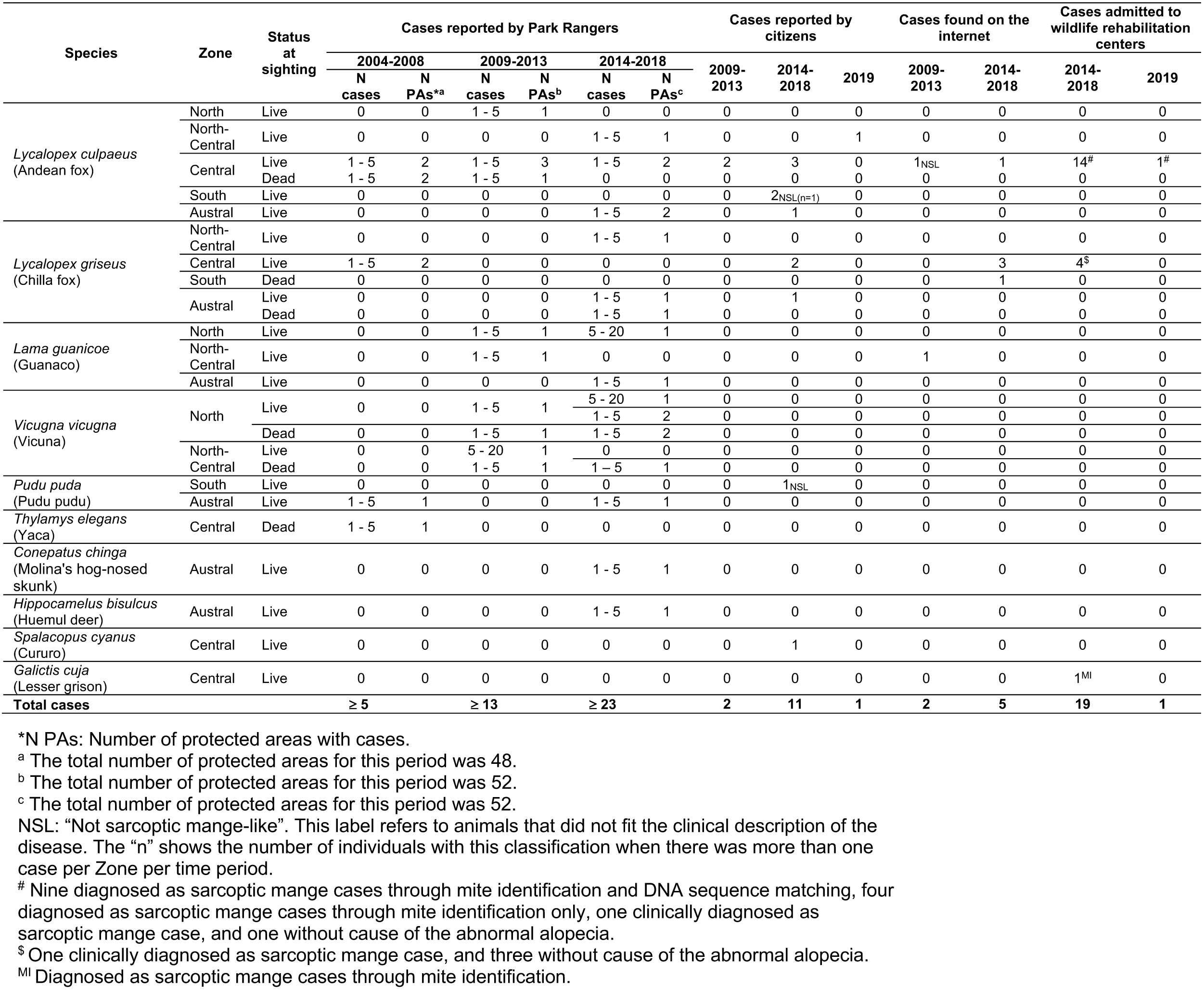
Number of wild mammals with abnormal alopecia within protected areas (park rangers) and outside protected areas (citizen reports; digital news outlets, social media, and others [internet]; and wildlife rehabilitation centers) per species, per Zone, and per period of time. In the case of protected areas, the number of cases per species, per Zone, and per period of time is also provided.

### b) Description of abnormal alopecia in wild mammals outside of Chilean protected areas

We received 17 web-based citizen reports during a 32-month period. Three reports were disregarded because pictures were absent or abnormal alopecia was arguable. The 14 animals in the valid reports were sighted between January 2013 and August 2019 from North-Central to Austral Chile. *Lycalopex sp*. were the group most commonly reported (85.71%; n=12; Table 2). These individuals showed alopecia (usually in the tail), hyperkeratosis, at least half of them were emaciated, and one had a rectal prolapse. Eleven of these twelve *Lycalopex* resembled sarcoptic mange infestation following previous descriptions (Bornstein et al., 2001; Niedringhaus et al., 2019; Pence and Ueckermann, 2002). Furthermore, a cururo (*Spalacopus cyanus*) and a pudu (*Pudu puda*) were also reported with evident hair loss. The overall appearance of the cururo suggests a probable sarcoptic mange case; conversely, the pudu does not resemble *S*. *scabiei* infestation. All photographs are available at 10.6084/m9.figshare.7797251.

We retrieved reports from the internet on four *L*. *griseus* and three *L*. *culpaeus* that were either found with abnormal alopecia (n=5) or reported as sarcoptic mange cases (n=2). These animals were observed in the Central and South Zones between 2012-2017, but three of them were clustered in 2016. The URLs for six of these cases are provided in S6. The remaining case corresponds to one previously confirmed *L*. *griseus* sarcoptic mange case found in the South Zone (Verdugo et al., 2016). One *L*. *culpaeus* case in the Central Zone during the second period was observed in a PA which reported cases of the same species for the same period and is not shown in Table 2 in the “Cases found on the internet” column, period “2014-2018” to avoid duplication. Six of the seven abnormally alopecic *Lycalopex* without confirmatory *S*. *scabiei* diagnosis resembled sarcoptic mange appearance (Bornstein et al., 2001; Niedringhaus et al., 2019; Pence and Ueckermann, 2002). One report of guanacos infested with sarcoptic mange in the North Central Zone for the period 2009-2013 was also retrieved (Zárate and Valencia, 2010).

Further, we extracted reports from previously collected data (Romero et al., 2019) on five *L*. *culpaeus*, four *L*. *griseus*, and a single lesser grison (*Galictis cuja*) with abnormal alopecia admitted to three different rehabilitation centers between 2013 and 2015. Of these, four *L*. *culpaeus*, one *L*. *griseus*, and the single *G*. *cuja* were confirmed *S*. *scabiei* cases (identification of the mite), while the records of the remaining individuals did not include information on the diagnosed cause of alopecia. These last ten animals came from the Central Zone (unknown Province). Finally, we gathered information from 12 additional *L*. *culpaeus* admitted to three rehabilitation centers between April 2016 and February 2019 that fulfilled our case definition. These animals presented hyperkeratosis, pyoderma, and emaciation and were all clinically diagnosed with sarcoptic mange. Sarcoptic mange was later confirmed by us in nine of them (see “Validity of alopecia to define a case of sarcoptic mange”). They were found in Central Chile and included two individuals observed in a PA which was positive for abnormally alopecic mammals according to the survey to park rangers during the same period and for the same species. These two individuals are not shown in Table 2 in the “Wildlife rehabilitation centers” column, period “2014-2018”, to avoid duplication. Fleas, ticks, and lesions caused by snare traps were additional findings reported in these cases.

Across these data sources, *Lycalopex* was the most common group found with abnormal alopecia (90.48%; 38/42). Based on the dates of observation and their location (excluding cases with unknown origin), these animals were all unique cases. The location of abnormally alopecic *Lycalopex* individuals (Provinces) that we were able to trace combining all data sources and that were observed between 2012-2019 (including the 2014-2018 period of the survey to park rangers) is shown in Fig 2. As stated previously, we acknowledge that these individuals were not necessarily present across the entire provincial territory, but we chose this scale to facilitate visualization.

**Fig 2.**
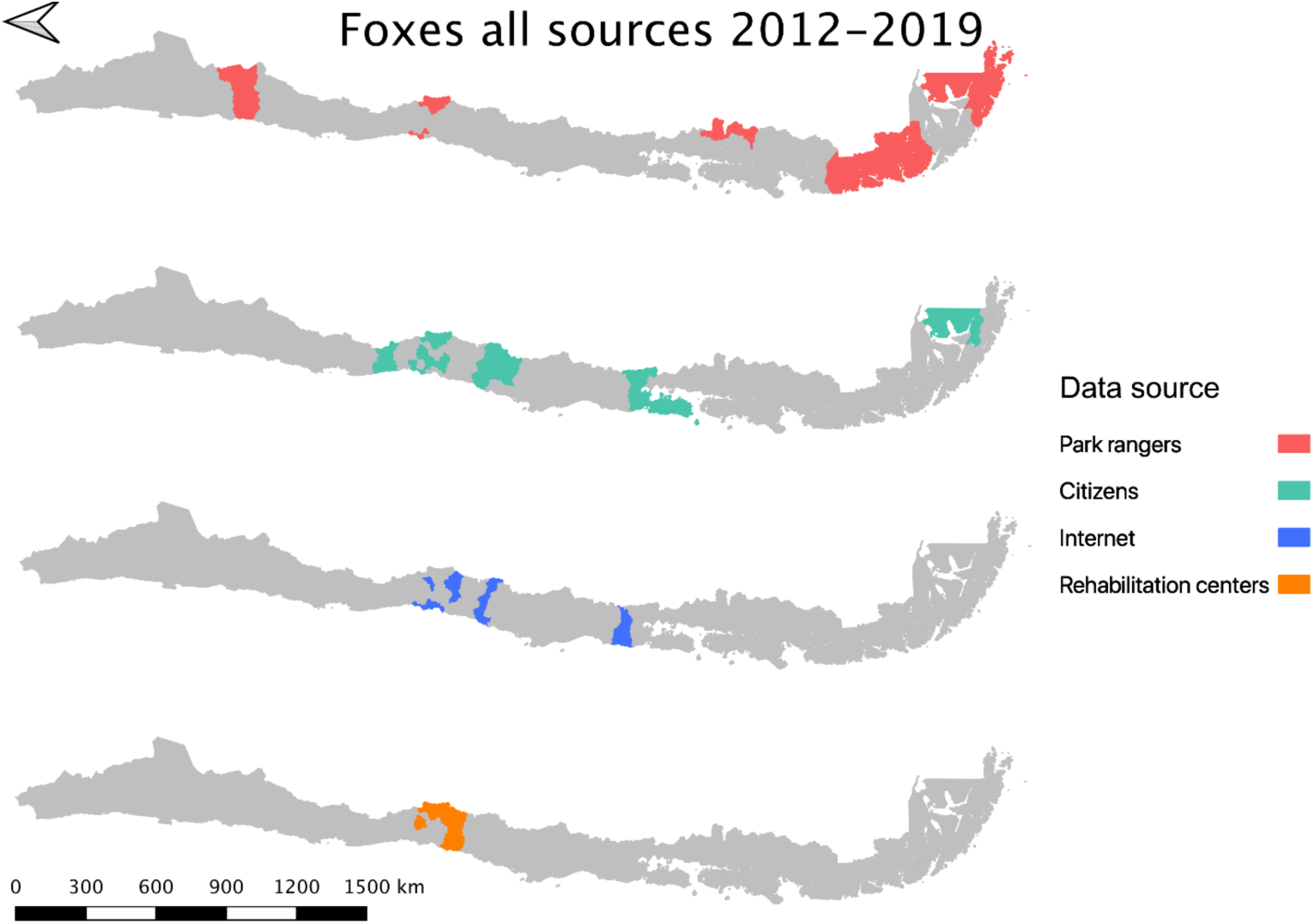
Location of abnormally alopecic *Lycalopex* foxes within and beyond protected areas between 2012-2019. The colored polygons are the Provinces that contain records of abnormally alopecic *Lycalopex* foxes (including confirmed sarcoptic mange cases) following the different data sources presented in Table 2. We present the information at this scale to facilitate visualization. This map also includes a previously reported case (Verdugo et al. 2016; orange) but does not include nine *Lycalopex* sp. admitted to rehabilitation centers according to Romero et al. (2019) whose Province of origin is unknown (but belonged to the Central Zone).

### c) Validity of alopecia to define a case of sarcoptic mange

Three rehabilitation centers provided skin samples from nine *L*. *culpaeus* that fulfilled our case definition (see above). Mites morphologically consistent with *S*. *scabiei* were isolated from all of them (S7). We conducted a molecular confirmation for all nine *L*. *culpaeus*. Sequencing in both directions of the internal transcribed spacer-2 (ITS-2) region produced a 404 bp sequence with 99.3% homology to *S*. *scabiei* (GenBank accession number A980753) for three of the nine *L*. *culpaeus* cases. We also sequenced in both directions the 16S rRNA region, producing a 331 bp fragment which rendered a 99.7% homology with *S*. *scabiei* (GenBank accession number MF083740) for four of the nine *L*. *culpaeus* cases. In summary, conventional PCR and sequencing performed on individual mites removed from the skin of infested foxes, molecularly confirmed this mite for three (ITS-2) and four (16S) of the nine cases. For the remaining two cases, sequences retrieved by PCR and sequencing were of insufficient quality, which may be due to fragmented DNA or inappropriate preservation methods at rescue centers of origin.

Given that all cases admitted to rehabilitation centers were *Lycalopex* foxes, we were only able to assess the validity of abnormal alopecia to track for *S*. *scabiei* infestation in this group. A binomial outcome of nine *S*. *scabiei* infested foxes (successes) out of nine with alopecia (*n*=9) has a probability to occur ≥ 0.5 if at least 0.923 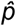 of the general population of alopecic foxes are, indeed, infested with this mite (Fig 3). The distance between the geographic origin of these nine foxes and their dates of admission suggest the independence of six foxes while the remaining three could be clustered. If this is correct and we consider this cluster as a single fox (for a total of 7 theoretical independent successes), then our conclusions of a very high likelihood of *S*. *scabiei* infestation in alopecic foxes remain valid (probability to observe infestation in all 7 foxes with alopecia is larger than 0.5 when 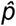 > 0.906).

**Fig 3.**
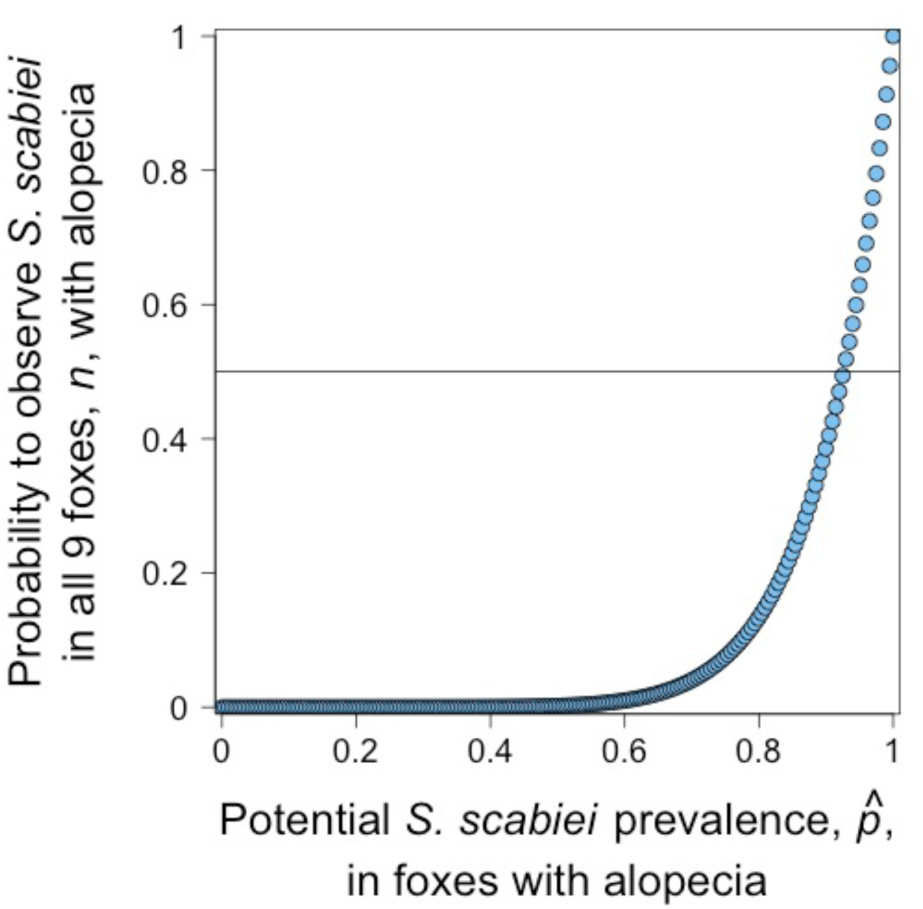
The probability of observing nine abnormally alopecic foxes (*n*) all of them infested with *Sarcoptes scabiei* under different potential sarcoptic mange prevalence, 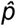. The horizontal line shows the 0.5 probability for this outcome.

## Discussion

Herein we collated data from several sources to describe, to the extent that current information allows, the presence of abnormally alopecic wild mammals in Chilean PAs over a 15-year period (2004–2018) and to summarize the location of affected animals beyond PAs. We observed a potential increment in the number of PAs in Chile in which wild mammals with abnormal alopecia have been sighted during the study period and a broad presence of cases across most of the Chilean territory in recent times. Data also support an increasing number of species observed with abnormal alopecia and a potential increment of cases within specific taxa. This suggested trend follows the current global emergence of sarcoptic mange in wildlife (Astorga et al., 2018; Niedringhaus et al., 2019).

*Lycalopex* foxes were the group most commonly reported with abnormal alopecia and were present across all data sources and time periods. The information from the Chilean National Park Service (CONAF) park rangers suggests that the geographical distribution of PAs presenting abnormally alopecic foxes has expanded over time. Between 2008-2018, these animals were observed in PAs located beyond Chile’s Central Zone where they were thought to be restricted (Cunazza and Diaz, 2014). Moreover, the incorporation of other sources of information, including citizen reports, allowed us to compile dozens of abnormally alopecic mammal reports from many parts of the country. Further, we confirmed *S*. *scabiei* in alopecic *L*. *culpaeus* from Central Chile, consistent with previous reports in an *L*. *griseus* from the southern part of the country and in other non-Chilean *Lycalopex* species (Deem et al., 2002; Verdugo et al., 2016). Through microscopic and molecular confirmatory testing, we showed that abnormal alopecia can be a good indicator of sarcoptic mange in *L*. *culpaeus*, similar to previous findings in red fox (*Vulpes vulpes*) with “mange-like lesions” (Nimmervoll et al., 2013) and that this disease is common in abnormally alopecic *Lycalopex* individuals.

South American camelids were the second most common group reported with abnormal alopecia. *S*. *scabiei*-infested South American camelids have been previously reported in PAs of the North and North Central Zones (Bonacic et al., 2014; López Jiménez, 2018) in agreement with our survey results. Park rangers support that abnormally alopecic vicuna and guanaco were found in a larger number of PAs and in a larger number of individuals during 2014-2018 compared to 2004-2008. However, whether this reflects an increasing trend is unclear. According to two published reports older than 2004 (and, therefore, not included in the internet data subset), sarcoptic mange in guanacos has been observed in the North Central Zone since 2002 (Zárate and Valencia, 2010) and before 2004 in the Austral Zone (Alvarado et al., 2004). Of note, pre-2004 sarcoptic mange reports in the Austral Zone refer to cases in central and southern Tierra del Fuego Island (Alvarado et al., 2004). No PA was established in central Tierra del Fuego until 2013; therefore, earlier South American camelid cases in the Austral Zone may have been under-reported in earlier years. For comparison, a distribution of South American camelid cases including all data sources and published pre-2004 data is shown in S8. Nevertheless, park ranger data presented here suggest a potential southward spread of abnormally alopecic South American camelids in the North Zone and sarcoptic mange in this taxon has been increasingly described in recent times in Peru, Bolivia, and Argentina (Castillo, 2018).

We also found cases affecting other species, such as a *Galictus cuja* diagnosed as a sarcoptic mange case through mite identification in a wildlife rehabilitation center. To our knowledge this is the first report of a lesser grison infested with *S*. *scabiei*, although sarcoptic mange has been previously reported in other mustelids (Mörner, 1992; Phillips et al., 1987).

Our results tend to support that abnormally alopecic wild mammals are observed more frequently, that these cases belong to a larger number of species over time, and that *S*. *scabiei* is a parsimonious explanation for some of the cases observed. Although the search for alopecic mammals through different detection methods, such as questionnaires, carcass collection, and camera traps have been useful to study sarcoptic mange in wildlife (Carricondo-Sanchez et al., 2017; Chen et al., 2012; Pisano et al., 2019; Soulsbury et al., 2007) these data sources are not without limitations. In our case, it is possible that self-selection bias occurred in the rangers that responded to the survey and that those rangers in PAs with alopecic mammals were more likely to participate. Nevertheless, if this were true, the proportion of affected PAs would be biased upwards, but the suggested temporal and spatial trends would remain. We also relied on historical records to avoid recall bias of alopecic mammal sightings by park rangers; however, the “surveillance” efforts and storage of this information could be better now than 20 years ago. We were informed by CONAF that differences in surveillance efforts and data recording were highly unlikely for the last two 5-year periods (2009-2013 and 2014-2018) when a potential increment in positive PAs was observed.

We advertised the web-platform to report alopecic wild mammals by several means and its value is justified in the recovery of alopecic cases that would otherwise not have been recorded. However, reporting may have been constrained by the visibility of alopecic animals or the degree of public awareness. This first experience suggests that web-based citizen reporting seems to be more useful for wild species present in easily accessed human-modified landscapes such as *Lycalopex* foxes versus South American camelids (or other taxa). This could explain the reports mostly belonging to the former group but none of the latter.

Further, we assessed the ability of abnormal alopecia to establish a suspect sarcoptic mange case by testing abnormally alopecic wild mammals for *S*. *scabiei*. We only had access to abnormally alopecic foxes, specifically, *L*. *culpaeus* that were admitted to wildlife rehabilitation centers; therefore, we could only assess the validity of this clinical sign to track sarcoptic mange in this species. This is relevant because the use of alopecia as a proxy to define a sarcoptic mange case in wildlife can be misleading (Valldeperes et al., 2019). We did not test free-ranging abnormal alopecic individuals of this species either. It is possible that *S*. *scabiei*-infested foxes are more likely to be captured and admitted into rehabilitation centers compared to other conditions leading to abnormal alopecia. If so, then our results may not apply to the general “abnormally alopecic fox” population. Nonetheless, although the number of foxes used to assess the validity of alopecia is not large, our results clearly show the need for a high “true prevalence” of *S*. *scabiei* to detect 9 out of 9 foxes infested. Park rangers (the only source of cases in our data for South American camelids) are most likely capable of distinguishing normal molting from abnormal alopecia. The external appearance of confirmed South American camelid sarcoptic mange cases from a recent outbreak in Argentina matches the observations reported by CONAF, providing more support for *S*. *scabiei* infestation in these animals (S5; Ferreyra et al., 2020).

Finally, we cannot claim that all abnormally alopecic mammals were infested with *S*. *scabiei*. Indeed, the reported huemul in the Austral Zone is probably an individual affected by *C*. *pseudotuberculosis* (Morales et al., 2017).

The data limitations mentioned above result from the fact that Chile, as is the case in all South American nations, does not have a National Wildlife Health Surveillance Program (Pinto et al., 2008). The lack of such a program hinders the collection of representative data on disease and etiologies in native mammal populations; our understanding of the epidemic or endemic nature of diseases, seasonal trends, and eventual populations impacts; and, most importantly, the response and management of health-related and potentially conservation-threatening issues. These threats include pathogens that have recently been detected in Chilean wildlife involving endemic endangered species. For example, caseous lymphadenopathy, echinococcosis, and a parapoxvirus-associated foot disease in the endangered huemul deer (Hernández et al., 2019; Morales et al., 2017; Vila et al., 2019); and Chytridiomycosis in the endangered Darwin’s frogs (*Rhinoderma* spp.; Soto-Azat et al., 2013). Moreover, several other pathogens have been sporadically detected in Chilean wildlife within PAs (Cunazza and Diaz, 2014; López Jiménez, 2018) whose status, trend, and relevance remain unknown.

Considering that several reports we collated correspond to confirmed and probable sarcoptic mange, its etiology, the mite *S*. *scabiei*, is a multi-host pathogen (Bornstein et al., 2001; Pence and Ueckermann, 2002), whose transmission may behave in a frequency-dependent manner (Devenish-Nelson et al., 2014), and infested hosts present a long infectious period. These traits have been associated with easier disease spread and disease-mediated extinction (Cross et al., 2005; De Castro and Bolker, 2005). Indeed, this mite is continuously reported in new species (Gonzalez-Astudillo et al., 2018), it can devastate populations of wild species (Martin et al., 2017) and it is currently considered a global emerging threat for wildlife conservation (Astorga et al., 2018). Further, *S*. *scabiei* may interact with other conservation threats or it could generate fragmented populations susceptible to stochastic events.

*Lycalopex* is a Neotropic endemic genus that belongs to a taxonomic family with worldwide records of *S*. *scabiei*-driven extirpations or near extirpations of common and endangered species (Henriksen et al., 1993; León-Vizcaíno et al., 1999; Lindström, 1991; Pence and Ueckermann, 2002; Soulsbury et al., 2007; Uraguchi et al., 2014). In Chile, *L*. *culpaeus* and *L*. *griseus* fulfill the description of the first group, while the endemic and endangered Darwin’s fox (*Lycalopex fulvipes*) and the endemic and vulnerable Fuegian culpeo (*L*. *c*. *lycoides*), certainly belong to the second category (Ministerio del Medio Ambiente, 2011; Silva-Rodríguez et al., 2016). These two endangered foxes would be highly at risk from any lethal disease outbreak. Of concern, mite-associated morbidity (*Otodectes cynotis*) has recently been reported in these two fox species (Briceño et al., 2019).

In the case of South American camelids, sarcoptic mange is considered a main threat for this group in Chile (Bonacic et al., 2014; Vargas et al., 2016; Zárate and Valencia, 2010). Moreover, an ongoing sarcoptic mange outbreak has nearly caused the local extinction of vicuna and guanaco in San Guillermo National Park in Argentina (Ferreyra et al., 2020), revealing the potential consequences of *S*. *scabiei* in these species. In addition to this infectious threat, the guanaco could become extinct in three of the five countries where it is currently present, projections for the conservation of South American camelids in the Chilean North Central Zone are not optimistic, and Chilean coastal populations have rapidly declined (Bonacic et al., 2014; González and Acebes, 2016; Vargas et al., 2016; Zárate and Valencia, 2010).

Despite being previously recognized in Chile and representing a potential threat for Chilean wildlife, data on *S*. *scabiei* in *Lycalopex sp*. and South American camelids are remarkably precarious and this is the first attempt to describe the status of this disease nationwide. Our findings, the potential consequences of this disease, and the absence of robust *S*. *scabiei* data further reinforce the need for a National Wildlife Health Surveillance Program.

Furthermore, it is anticipated that disease emergence will occur more often in wild species globally as a result of the human-mediated movement of pathogens, parasites, and domestic hosts, and concurrent changes in the environment including habitat destruction and the climate emergency (Daszak et al., 2000). These drivers are occurring in Chile. For example, 30% of native temperate forest was lost during the last three decades (Zamorano-Elgueta et al., 2015) and 69% of mammals were already threatened by habitat destruction in Central Chile twenty years ago (Simonetti, 1999). Consequently, new disease events in Chilean native species should be expected in the upcoming years.

Based on our previous arguments we encourage the urgent convening of an intersectoral working group of private stakeholders, academia, and public agencies with the goal of establishing a National Wildlife Health Surveillance Program in Chile. Existing uncertainties with respect to sarcoptic mange levels and trends and the potential threat posed by this disease justify its prompt launch and provides an opportunity for capacity-building and troubleshooting during start-up. Such a program should fulfill the attributes, purposes, and roles recently proposed by experts (Stephen et al., 2018) and should engage domestic animal and human health sectors as well as conservation and biodiversity constituents through a One Health approach. Government-led wildlife health programs have had remarkable positive influences on the conservation of species as well as on public health (Brand, 2013; Kuiken et al., 2005; Reif, 2011). Although there is an incipient valuable pilot effort lead by the Chilean National Park Service to monitor diseases, it only covers some PAs and several components need to be expanded or substantially improved to generate a functional country-level Wildlife Health Program (Aizman et al., 2016; Cunazza et al., 2015). Nations failing to properly prepare to confront pathogen threats in wildlife may expect to experience catastrophic outcomes for conservation as described worldwide (De Castro and Bolker, 2005). Moreover, governments should strive towards fulfilling their responsibilities to meet societal needs and obligations in their role as keepers of a public good.

## Conclusions

Abnormally alopecic wild mammals seem to have become more common in Chile and observed in an increasing area of the country over a 15-year period, between 2004-2018. Sarcoptic mange is a likely cause for this disease at least in the genus most commonly reported with alopecia, *Lycalopex*, and in South American camelids. Based on these findings and the catastrophic consequences that sarcoptic mange has had worldwide, we encourage the establishment of a National Wildlife Health Surveillance Program in Chile. Such a program is also justified by the recent detection of pathogens causing morbidity and mortality in native species, including the iconic Huemul deer, classified as endangered in the IUCN’s Red List.

## Acknowledgments

We appreciate the support from Miguel Díaz and Gabriela López from CONAF, as well as all the park rangers who answered our survey. We thank all the citizens who took the time to report and provide us with pictures and information. We are also thankful to Rayén Díaz for the *Sarcoptes scabiei* pictures and Violeta Barrera and Jose Caro, whose comments improved the survey. We are grateful to Parque Safari, especially Diego Peñaloza; Unidad de Rehabilitación de Fauna Silvestre UFAS, especially Nicole Sallaberry; Centro de Rescate y Rehabilitación Hospital Clínico Veterinario Universidad Santo Tomás, Viña del Mar; and Zoológico Nacional de Chile, especially Andrea Caiozzi, Andrés Ramírez, and Marisol Torregrosa, for their valuable support in sample and data collection. We kindly appreciate the constructive input of three reviewers.

