## Supplementary material for "Sarcoptic mange; an emerging threat to Chilean wild mammals?": S1, S2, S3, S4, S5, S6, S7, S8

### Supporting Information 1

**Survey applied to the rangers in National Protected Areas in Spanish and English.**

### Casos sospechosos de sarna en areas silvestres protegidas del Estado

**Muchas gracias por su interés en participar de esta encuesta y aportar con su información para poder comprender que está ocurriendo con los avistamientos de mamíferos con pérdida de pelo a nivel nacional. En Chile desde hace algunos años se ha capturado zorros con pérdida de pelo que han resultado positivos a sarna sarcóptica. La sarna sarcóptica es una enfermedad infecciosa causada por el ácaro microscópico *Sarcoptes scabiei*, que habita en la piel de los animales infectados. Aquí pone huevos, se alimenta, defeca, y genera inflamación e infección bacteriana secundaria de la piel. Esta enfermedad causa evidente pérdida de pelo y puede ocasionar la muerte del individuo infectado. Pero no sólo ocurre en zorros. De hecho este parásito ha sido detectado en más de 100 especies de mamíferos, y se considera una enfermedad emergente en fauna silvestre alrededor del mundo. Este ácaro es capaz de generar impactos poblacionales importantes. Por ejemplo, en Europa se ha cuantificado disminuciones poblacionales de hasta un 95% de zorros rojos, y se han registrado extinciones locales a causa de esta enfermedad (zorro rojo y Wombats en Australia; si quiere saber más de esta enfermedad haga click aquí). Por lo tanto, dado que la sarna sarcóptica puede ser relevante para la conservación de la biodiversidad,**
**queremos desarrollar una descripción inicial de esta enfermedad a escala nacional para saber, por ejemplo, dónde se ha visto, en qué especies, etc. Su participación es muy importante para cumplir con este objetivo. Por favor, es vital que sólo se responda una encuesta por Área Protegida. Es decir, si usted responde esta encuesta para caracterizar el Área Protegida donde trabaja, entonces ningún colega debe contestar esta encuesta para la misma Área Protegida. Si usted ha trabajado menos de 15 años en el Área Protegida para la cual va a responder la encuesta, por favor recopile la información necesaria para contestar correctamente las preguntas que se refieren a periodos anteriores a su llegada a dicha Área Protegida. En caso de dudas por favor escriba a .**

1. Por favor adjunte la información requerida

**Área protegida en la que se desempeña** __________

**Comuna donde se ubica el Área Protegida** __________

**Región donde se ubica el Área Protegida** __________

**Desde cuando trabaja en dicha Área protegida** __________

2. En el área protegida donde usted trabaja, se han avistadomamíferos vivos o cadáveres con pérdida de pelo en los últimos 15 años que se hayan visto como en en las imágenes mostradas abajo? En caso de que SI se hayan avistado este tipo de animales, por favor marque todos los períodos que correspondan (puede marcar más de una respuesta). En caso de que NO se hayan avistado animales con pérdida de pelo en los últimos 15 años por favor haga click en la opción "No se han avistado" y la encuesta llegará a su fin.

- ☐ Entre 15 y 10 años atrás
- ☐ En los últimos 5 años
- ☐ Entre 10 y 5 años atrás
- ☐ No se han avistado

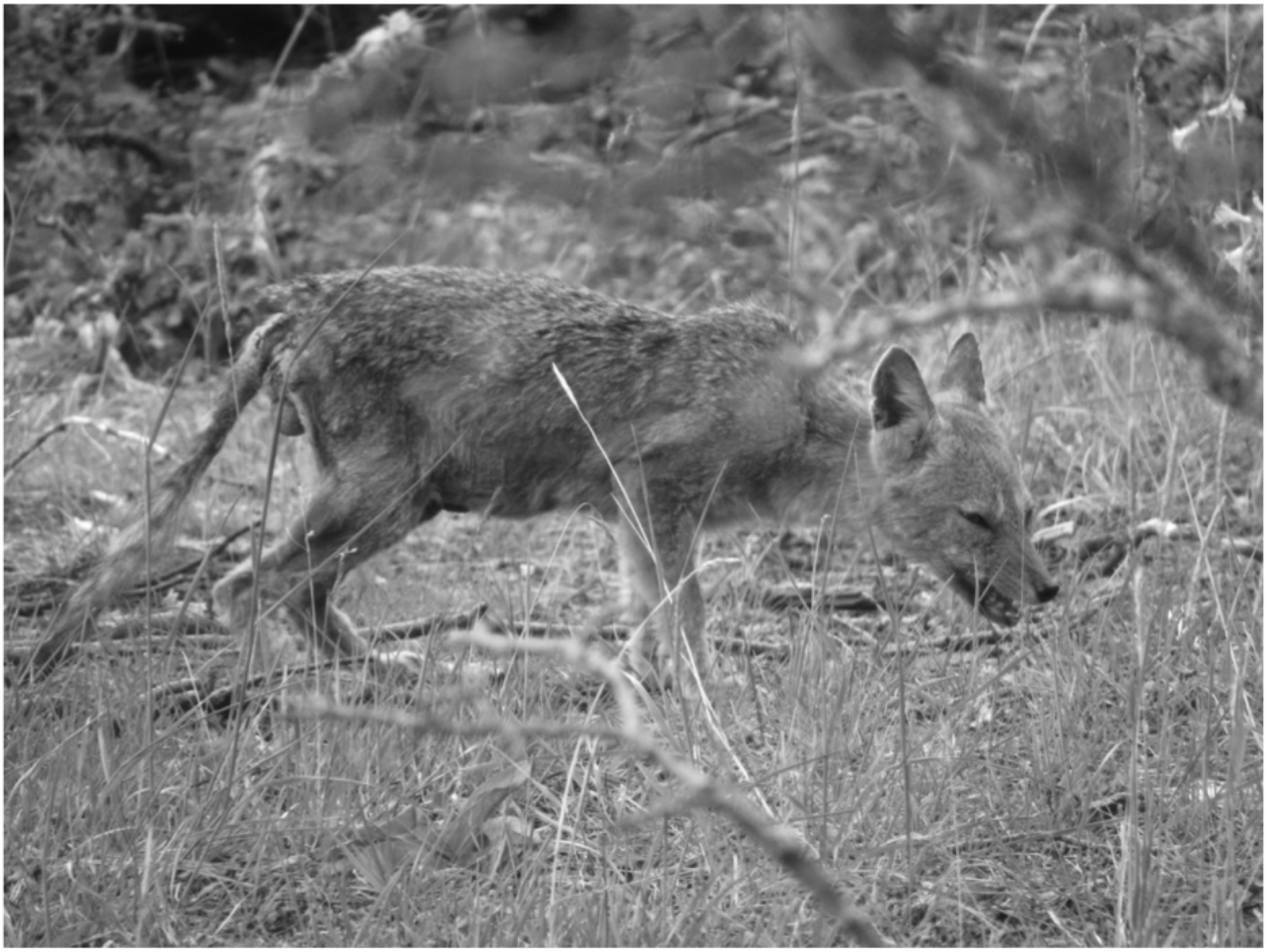

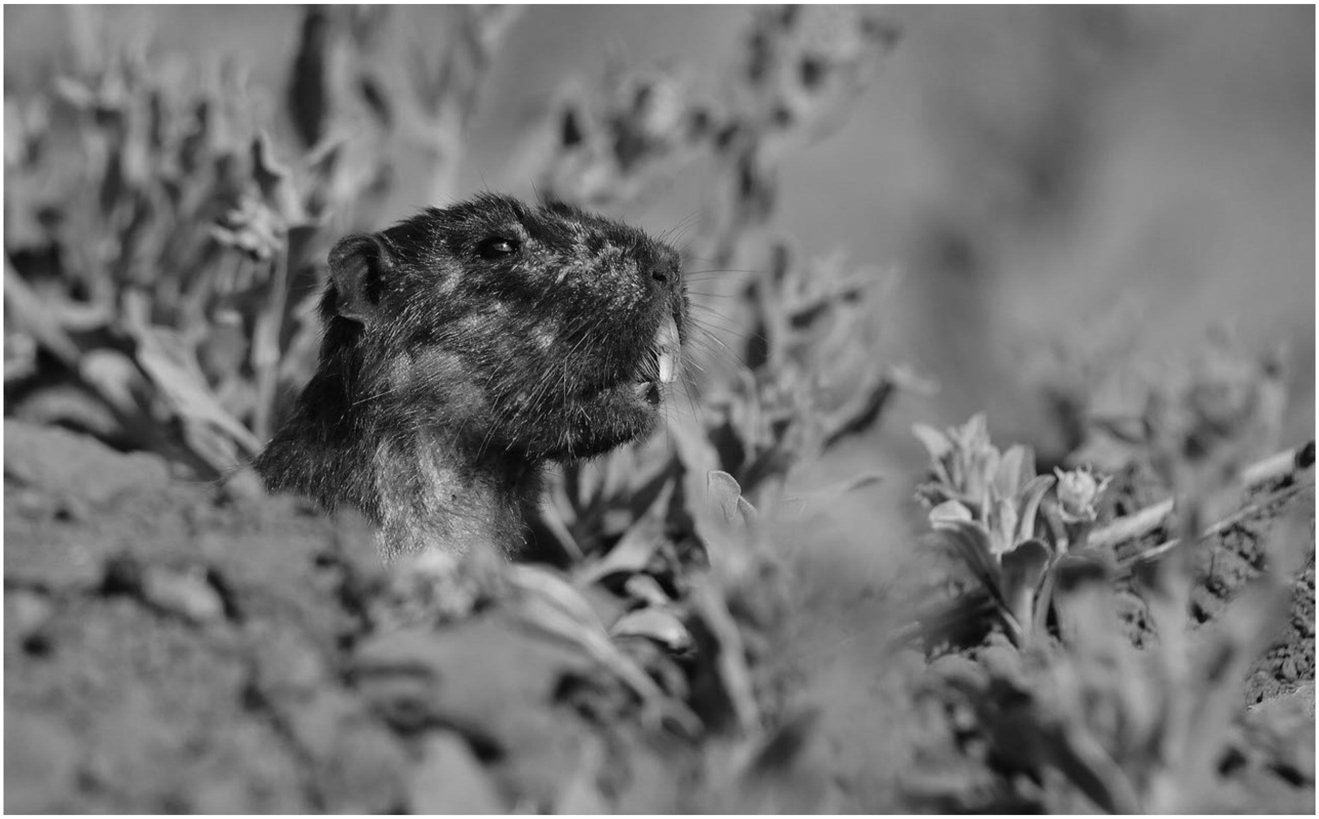

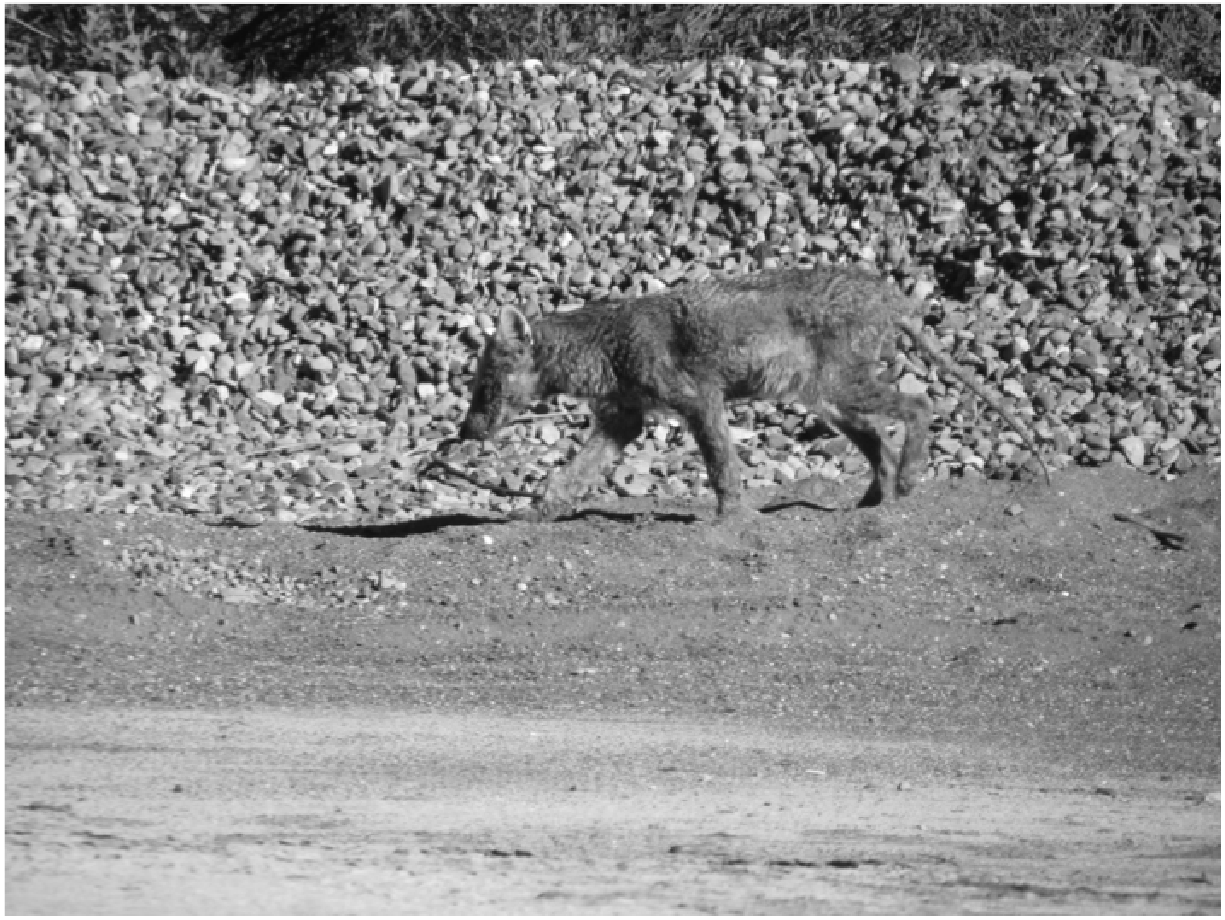

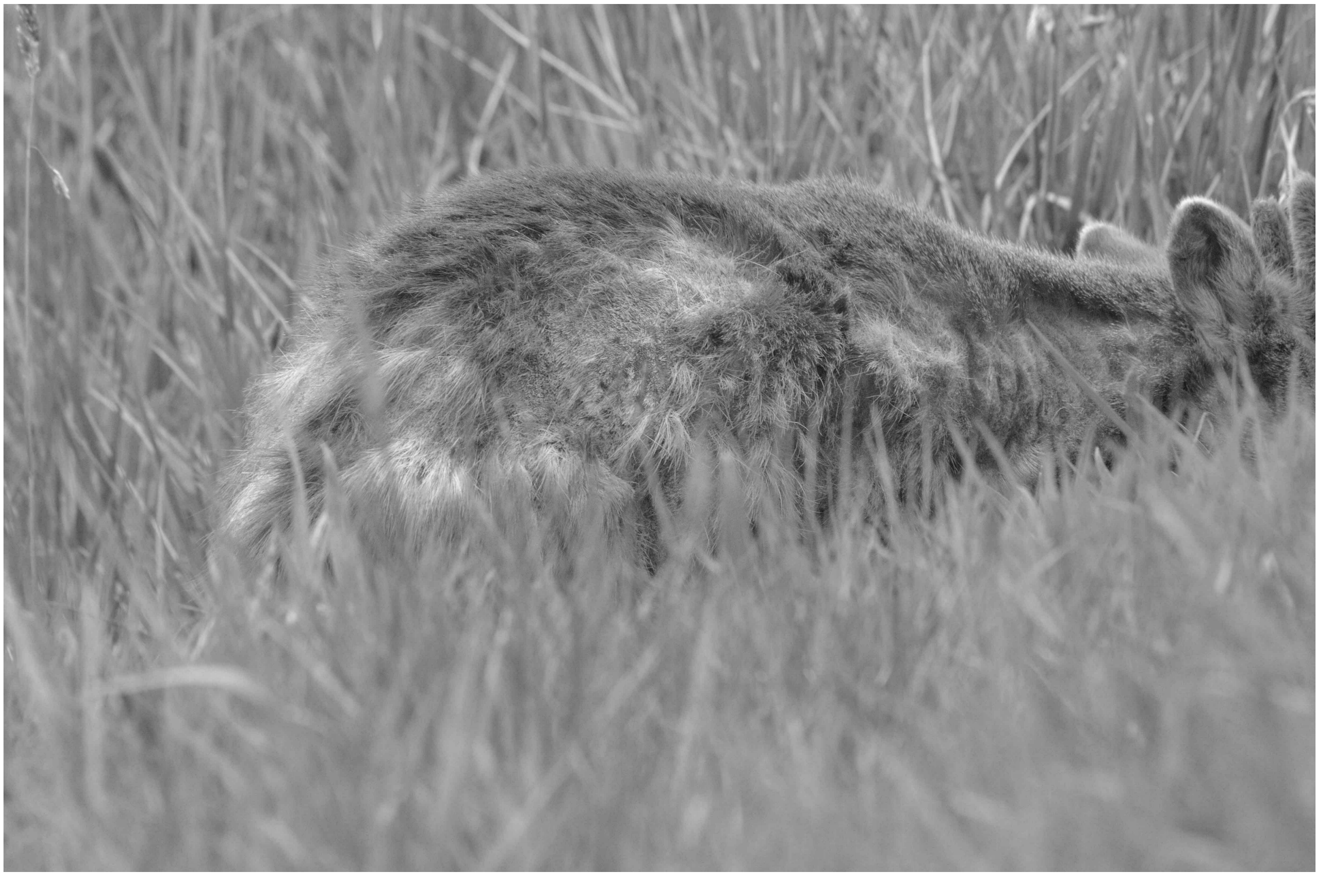

3. Esta pregunta se refiere a animales vivos. Si en el Área Protegida donde usted trabaja se han avistado mamíferos con pérdida de pelo, por favor marque todos los períodos en los que se encontraron a estos mamíferos (entre 15 y 10 años atrás, entre 15 y 10 años atrás y en los últimos 5 años). En caso de que no se hayan avistado mamíferos con pérdida de pelo en ningún período marque "No se han encontrado" y automáticamente pasará a la pregunta 5.

- ☐ Entre 15 y 10 años atrás
- ☐ Entre 10 y 5 años atrás
- ☐ En los últimos 5 años
- ☐ No se han encontrado

4. Esta pregunta se refiere a animales vivos. Por favor indique el número de animales encontrados con pérdida de pelo por período de tiempo. Si hay roedores en la lista, por favor indique la especie al final de la tabla. Si ha observado otra especie, por favor indíquela al final de la tabla. Para las especies en las que no se ha observado animales con pérdida de pelo en ningún período, por favor no marque ninguna opción.

| | Entre 15 y 10 años atrás | Entre 10 y 5 años atrás | En los últimos 5 años |
|---|---|---|---|
| Zorro chilla | | | |
| Zorro culpeo | | | |
| Zorro de Darwin | | | |
| Perro doméstico | | | |

| | Entre 15 y 10 años atrás | Entre 10 y 5 años atrás | En los últimos 5 años |
|---|---|---|---|
| Guanaco | | | |
| Vicuña | | | |
| Taruca | | | |
| Puma | | | |
| Gato Andino | | | |
| Gato Colo Colo | | | |
| Gato de Geoffroy | | | |
| Güiña | | | |
| Quique | | | |
| Chingue | | | |
| Pudú | | | |
| Huemul | | | |
| Jabalí | | | |
| Ciervo rojo | | | |
| Conejo/Liebre | | | |
| Visón | | | |
| Vaca | | | |
| Caballo | | | |
| Burro | | | |
| Cabra | | | |
| Oveja | | | |
| Roedor 1 | | | |
| Roedor 2 | | | |
| Roedor 3 | | | |
| Llaca | | | |

| | Entre 15 y 10 años atrás | Entre 10 y 5 años atrás | En los últimos 5 años |
|---|---|---|---|
| Comadrejita trompuda | | | |
| Quirquincho | | | |
| Huroncito patagónico | | | |
| Monito del monte | | | |
| Otra especie | | | |

Roedor 1, Roedor 2, Roedor 3, Otra especie

5. Esta pregunta se refiere a animales encontrados muertos. En el Área Protegida donde usted trabaja, se han encontrado cadáveres completos o partes de cadáveres de mamíferos que tuvieran pérdida de pelo no causada por la descomposición? Por favor marque todos los períodos en los que se encontraron cadáveres con pérdida de pelo (Entre 15 y 10 años, Entre 10 y 5 años, En los últimos 5 años). En caso de que no se hayan encontrado cadáveres de mamíferos con pérdida de pelo en ningún período marque "No se han encontrado" y automáticamente pasará a la pregunta 7.

- ☐ Entre 15 y 10 años atrás
- ☐ Entre 10 y 5 años atrás
- ☐ En los últimos 5 años
- ☐ No se han encontrado

6. Esta pregunta se refiere a animales encontrados muertos. Por favor indique el número de cadáveres encontrados con pérdida de pelo por período de tiempo. Si hay roedores en la lista, por favor indique la especie la final de la tabla. Si ha observado otra especie, por favor indíquela al final de la tabla. Para las especies en las que no se ha encontrado cadáveres con pérdida de pelo en ningún período por favor no marque ninguna opción.

| | Entre 15 y 10 años atrás | Entre 10 y 5 años atrás | En los últimos 5 años |
|---|---|---|---|
| Zorro chilla | | | |
| Zorro culpeo | | | |
| Zorro de Darwin | | | |
| Perro doméstico | | | |
| Guanaco | | | |
| Vicuña | | | |
| Taruca | | | |
| Puma | | | |
| Gato Andino | | | |

| | Entre 15 y 10 años atrás | Entre 10 y 5 años atrás | En los últimos 5 años |
|---|---|---|---|
| Gato Colo Colo | | | |
| Gato de Geoffroy | | | |
| Güiña | | | |
| Quique | | | |
| Chingue | | | |
| Pudú | | | |
| Huemul | | | |
| Jabalí | | | |
| Ciervo rojo | | | |
| Conejo/Liebre | | | |
| Visón | | | |
| Vaca | | | |
| Caballo | | | |
| Burro | | | |
| Cabra | | | |
| Oveja | | | |
| Roedor 1 | | | |
| Roedor 2 | | | |
| Roedor 3 | | | |
| Llaca | | | |
| Comadrejita trompuda | | | |
| Quirquincho | | | |
| Huroncito patagónico | | | |
| Monito del monte | | | |
| Otra especie | | | |

Roedor 1, Roedor 2, Roedor 3, Otra especie

7. Si tiene fotos de animales afectados o de cadáveres con pérdida de pelo, sería de utilidad que pudiese enviarlas a junto con la especie, lugar o Área protegida, y fecha de la fotografía.

- [ ] Sí tengo
- [ ] No tengo
- [ ] Sí tengo pero prefiero no compartirlas

8. Esta pregunta se refiere a animales vivos y muertos. En qué estación del año es más frecuente encontrar mamíferos con pérdida de pelo? Marque más de una estación si es necesario. En caso de no haber estacionalidad en el hallazgo de mamíferos con pérdida de pelo, por favor marque "No he visto más casos en una estación determinada".

- [ ] Otoño
- [ ] Invierno
- [ ] Primavera
- [ ] Verano
- [ ] No he visto más casos en una estación determinada

9. En general, en qué estación del año se realizan exploraciones del Área Protegida que abarquen mayor superficie y que sean más frecuentes. Marque más de una estación si es necesario. En caso de que las exploraciones sean similares durante todo el año, por favor marque "Es constante".

- ○ Otoño
- ○ Invierno
- ○ Primavera
- ○ Verano
- ○ Es constante

10. Si lo considera necesario, por favor utilice este espacio para dar otros detalles respecto a mamíferos con pérdida de pelo en el Área protegida donde trabaja que considere relevantes

Thank you very much for your willingness to participate in this survey and provide information aiming to understand what is going on with the sightings of mammals with hair loss nationwide. Since some years ago, foxes with hair loss positive to sarcoptic mange have been captured in Chile. Sarcoptic mange is an infectious disease caused by the mite *Sarcoptes scabiei,* which lives in the skin of the infested animals. Here, these mites lay eggs, feed, defecate, and causes inflammation leading to secondary infection of the skin. This disease causes evident hair loss and may lead to death. But this disease does not only occur in foxes. In fact, this parasite has been detected in more than a hundred mammal species and it is considered an emergent disease in wildlife globally. This mite can cause important population impacts. For example, 95% population reduction has been quantified in red foxes in Europe and local extinctions have been recorded in Europe due to this disease (red foxes and wombats in Australia; if you want to learn more about this disease please click here). Therefore, because sarcoptic mange can be relevant for biodiversity conservation, we want to develop an initial description of this disease nationwide in order to learn, for example, where it has been observed, in which species, etc. Your participation is very important to fulfill this objective. Please, it is extremely important that only a single survey is responded per Protected Area. This means that if you respond this survey to characterize the Protected Area where you work, then no other colleague should respond this survey for the same Protected Area. If you have worked less than 15 years in the Protected Area you are responding this survey for, please compile the records and information needed to answer the questions about past periods prior to your arrival to the corresponding Protected Area. If you have any doubts please contact us at.

1. **Please provide the information requested.**

Protected Area in which you currently work.

Commune where this Protected Area is located

Region where this Protected Area is located

When did you start working in this Protected Area?

2. **Have there been sightings of live or dead animals with hair loss in the last 15 years in the Protected Area you currently work that resemble the animals shown below?**

**If the answer is yes, please check all the appropriate periods. If the answer is no, please click in the option "They have not been observed" and the survey will be over.**

Between 15 -  10 years ago

Between 10 – 5 years ago

In the last 5 years

They have not been found

3. **This question refers to live animals. If mammals with hair loss have been observed in the Protected Area where you currently work, please check all the periods in which these mammals have been observed (between 15 – 10 years ago, between 10 – 5 years ago, and in the last 5 years). If no live mammal with hair loss has been observed in any period, check "They have not been observed" and you will be moved to question number 5.**

Between 15 -  10 years ago

Between 10 – 5 years ago

In the last 5 years

They have not been observed

4. **This question refers to live animals. Please indicate the number of animals observed with hair loss per time period. If there are rodents in the list, please name the specific species at the end of the following table. For those species without any individuals with hair loss, please do not provide any option.**

| | Between 15 - 10 years ago | Between 10 – 5 years ago | In the last 5 years |
|---|---|---|---|
| Chilla fox | | | |
| Culpeo fox | | | |
| Darwin fox | | | |
| Domestic dog | | | |
| Guanaco | | | |
| Vicuna | | | |
| Taruca | | | |
| Puma | | | |
| Gato Andino | | | |
| Gato Colo Colo | | | |
| Gato de Geoffroy | | | |
| Guiña | | | |
| Quique | | | |
| Chingue | | | |
| Pudu | | | |
| Huemul | | | |
| Wild boar | | | |
| Red Deer | | | |
| Rabbit/Hare | | | |
| Mink | | | |
| Cow | | | |
| Horse | | | |
| Donkey | | | |

| |
|---|
| Goat |
| Sheep |
| Rodent 1 |
| Rodent 2 |
| Rodent 3 |
| Mouse opossum |
| Shrew opossum |
| Andean hairy armadillo |
| Patagonian weasel |
| Colocolo opossum |
| Another species |

**Rodent 1, Rodent 2, Rodent 3, another species:**

5. **This question refers to dead animals. Have full or partial mammal carcasses been found in the Protected Area you work showing hair loss unrelated to decomposition? Please check all the periods in which these carcasses were found (between 15 – 10 years ago, between 10 – 5 years ago, and in the last 5 years). If no carcass with hair loss has been observed in any period, check "They have not been observed" and you will be moved to question number 5.**

Between 15 - 10 years ago

Between 10 – 5 years ago

In the last 5 years

They have not been observed

6. **This question refers to dead animals. Please indicate the number of carcasses found with hair loss per time period. If there are rodents in the list, please name the specific species at the end of the following table. For those species without any carcasses with hair loss, please do not provide any option.**

| | Between 15 - 10 years ago | Between 10 – 5 years ago | In the last 5 years |
|---|---|---|---|
| Chilla fox | | | |
| Culpeo fox | | | |
| Darwin fox | | | |
| Domestic dog | | | |
| Guanaco | | | |
| Vicuna | | | |
| Taruca | | | |
| Puma | | | |
| Gato Andino | | | |
| Gato Colo Colo | | | |
| Gato de Geoffroy | | | |
| Guiña | | | |
| Quique | | | |
| Chingue | | | |
| Pudu | | | |
| Huemul | | | |
| Wild boar | | | |

| |
|---|
| Red Deer |
| Rabbit/Hare |
| Mink |
| Cow |
| Horse |
| Donkey |
| Goat |
| Sheep |
| Rodent 1 |
| Rodent 2 |
| Rodent 3 |
| Mouse opossum |
| Shrew opossum |
| Andean hairy armadillo |
| Patagonian weasel |
| Colocolo opossum |
| Another species |

Rodent 1, Rodent 2, Rodent 3, another species:

7. **If you have pictures of affected animals or carcasses with hair loss, it would be useful you could send them to us naming the species, location or Protected Area and date of the picture.**

I have

I do not have

I have but I prefer not to share them

8. **This question is related to live and dead animals. What season of the year is more frequent to find mammals with hair loss? Mark more than a season if necessary.**

Fall

Winter

Spring

Summer

I have not observed more cases in a specific season

9. **In general, what season of the year is the Protected Area more explored encompassing larger surface and with higher frequency? Mark more than one season if necessary. If the explorations are similar across the year please mark "they are constant".**

Fall

Winter

Spring

Summer

They are constant

10. **If you think is necessary please use this space to provide other details with respect to mammals with hair loss in the Protected Area where you work.**

### Supporting Information 2

**Flyer to create awareness of the website collecting information of cases of wild mammals with alopecia**

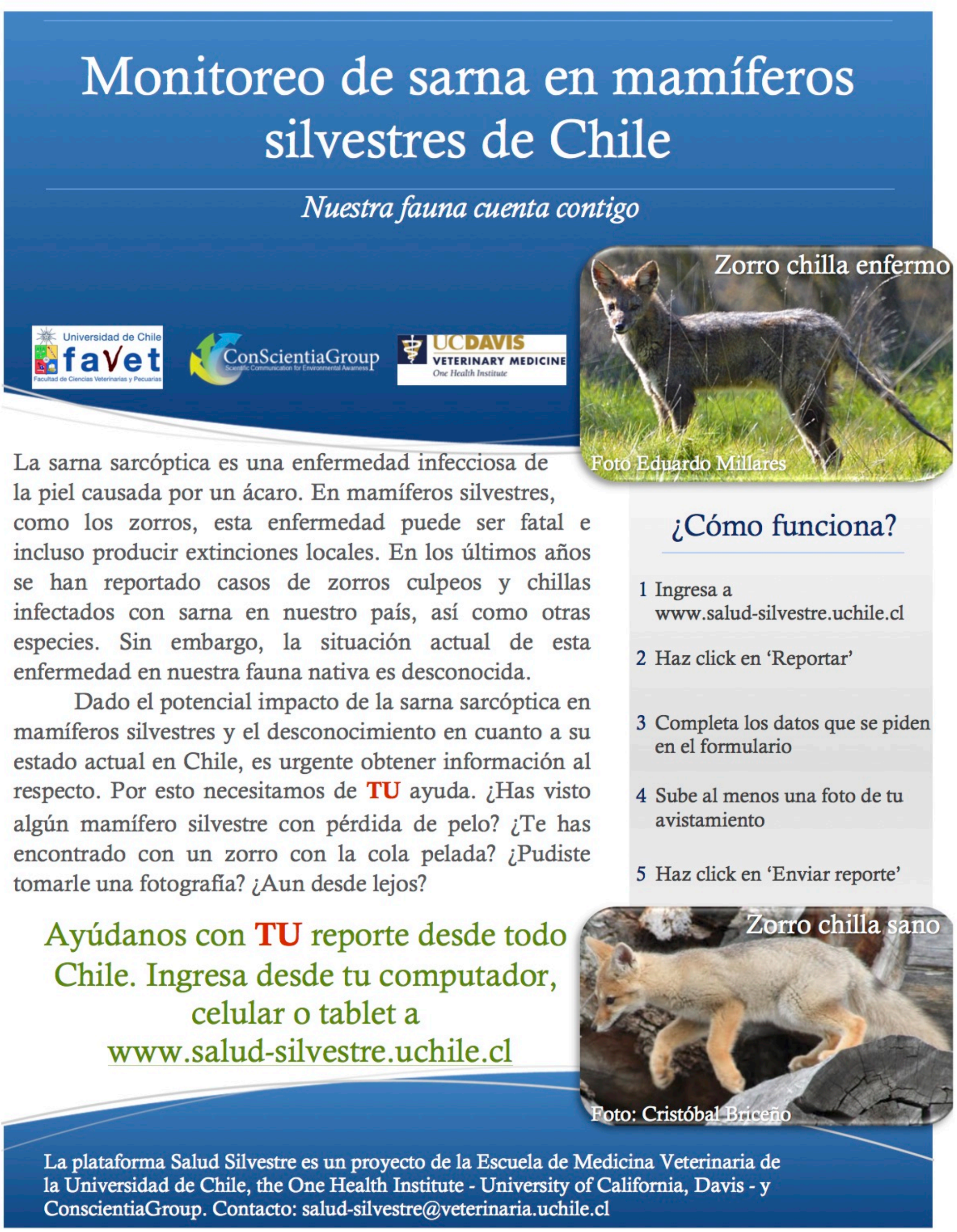

### Supporting Information 3

**Distribution of protected areas (orange) across Chile, per Zone (see S5), and the protected areas whose head ranger completed the survey (hashed areas).**

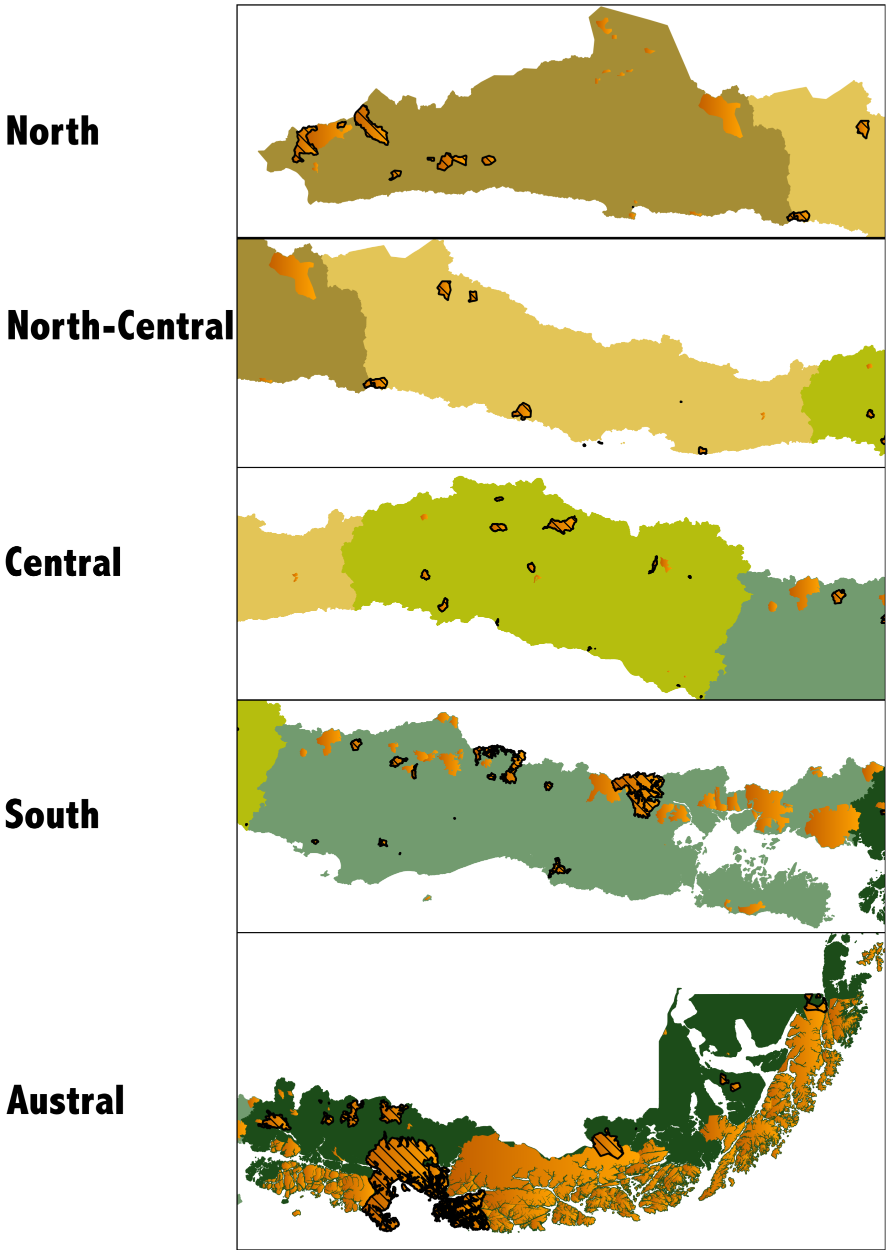

### Supporting Information 4

#### Alopecic wild mammals observed by CONAF park rangers

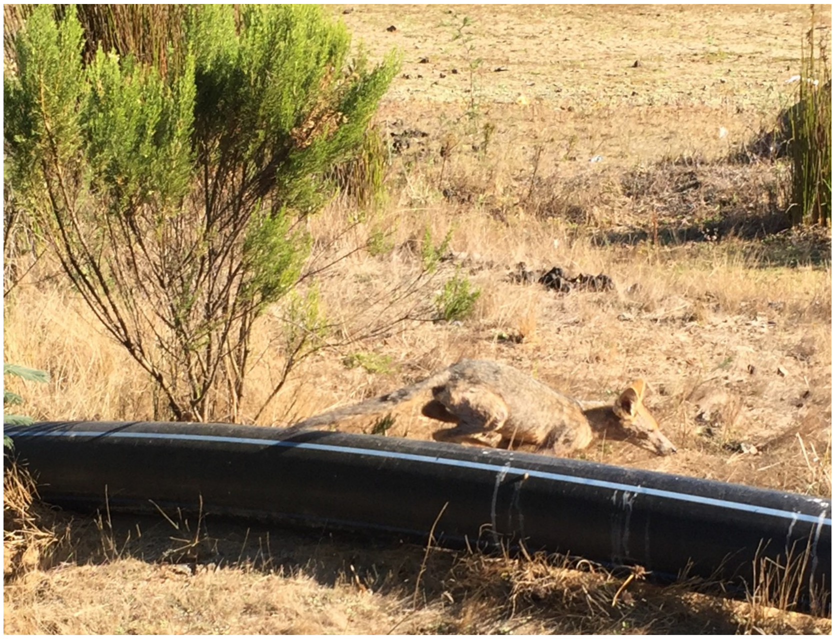

*Lycalopex culpaeus*

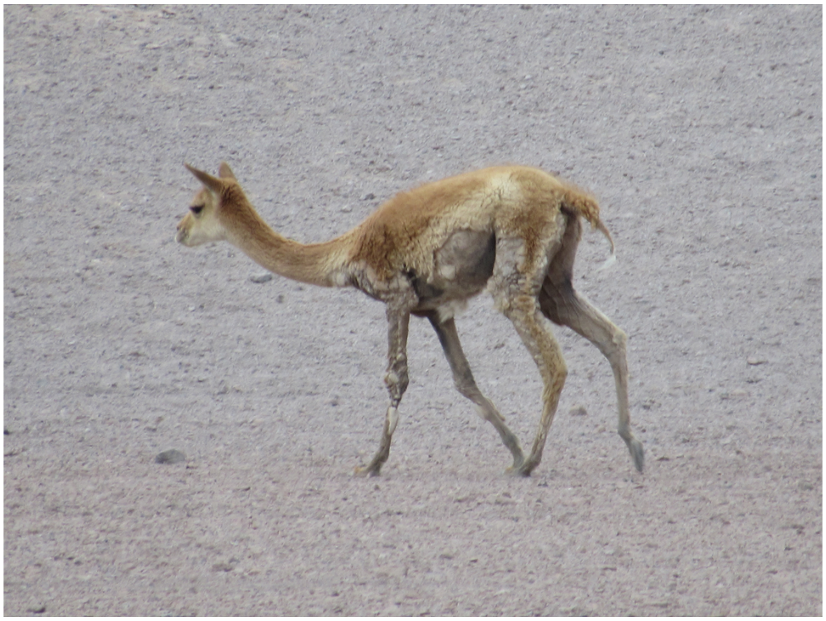

*Vicugna vicugna*

### Supporting Information 5

#### Geographical zones considered in the manuscript

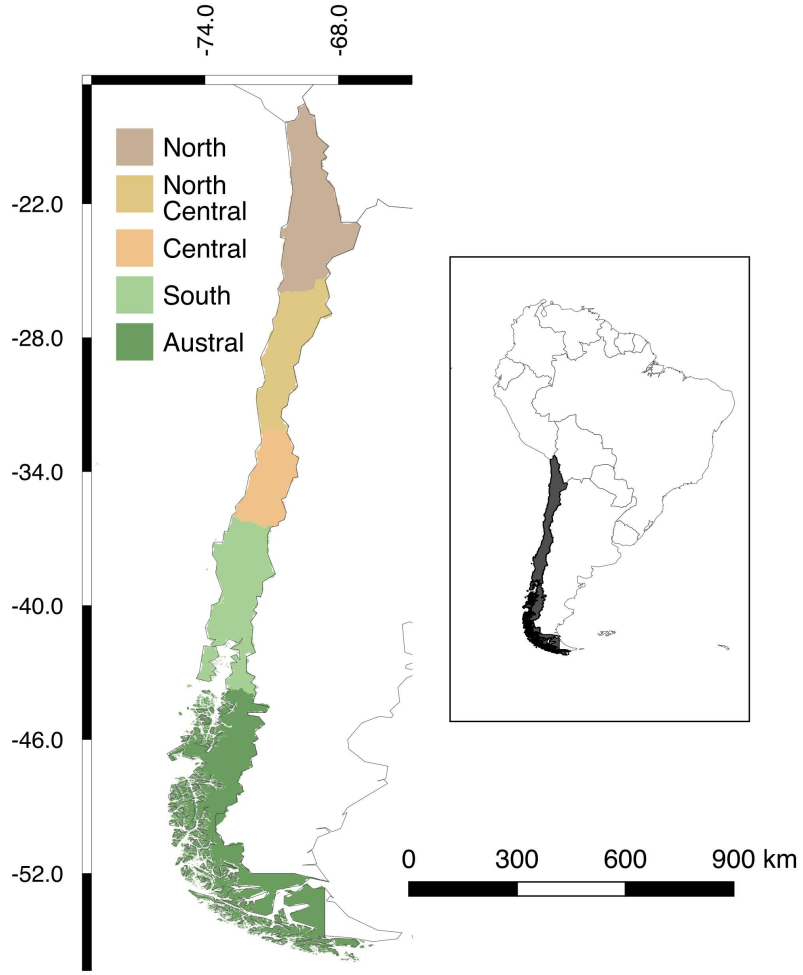

**Supporting Information 6**

**URL sarcoptic mange cases:**

- https://laderasur.com/articulo/cuando-la-sangre-tira-el-problema-de-los-perros-y-gatos-asilvestrados-con-la-fauna-nativa/

- https://www.flickr.com/photos/69758141@N03/6845820606

- https://www.soychile.cl/San-Antonio/Sociedad/2016/01/21/371171/La-impresionante-recuperacion-del-zorro-chilla-con-sarna-en-centro-de-rescate-de-San-Antonio.aspx

- https://twitter.com/informetierra/status/737529549634686976

- https://www.facebook.com/csv.sancarlosdeapoquindo/videos/vb.1518022025170688/1714204592219096/?type=2&theater

- https://twitter.com/Jo_infantev/status/899453925782409216

- http://intra.conaf.cl/descarga/planes-nacionales-de-conservacion-de-especies-fauna/?wpdmdl=9556&ind=5

### Supporting Information 7

***Sarcoptic scabiei* recovered from *Lycalopex sp*.**

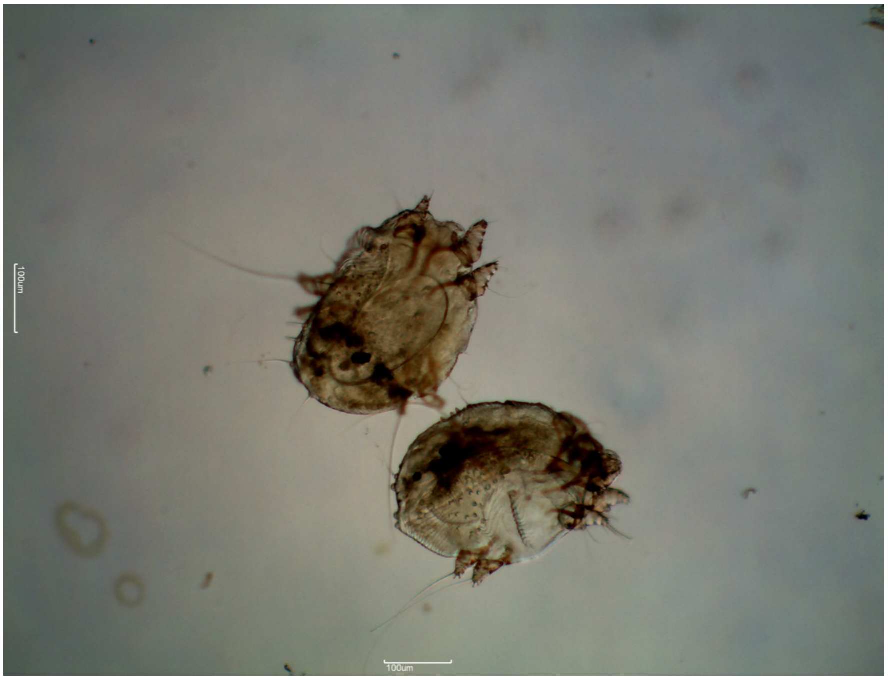

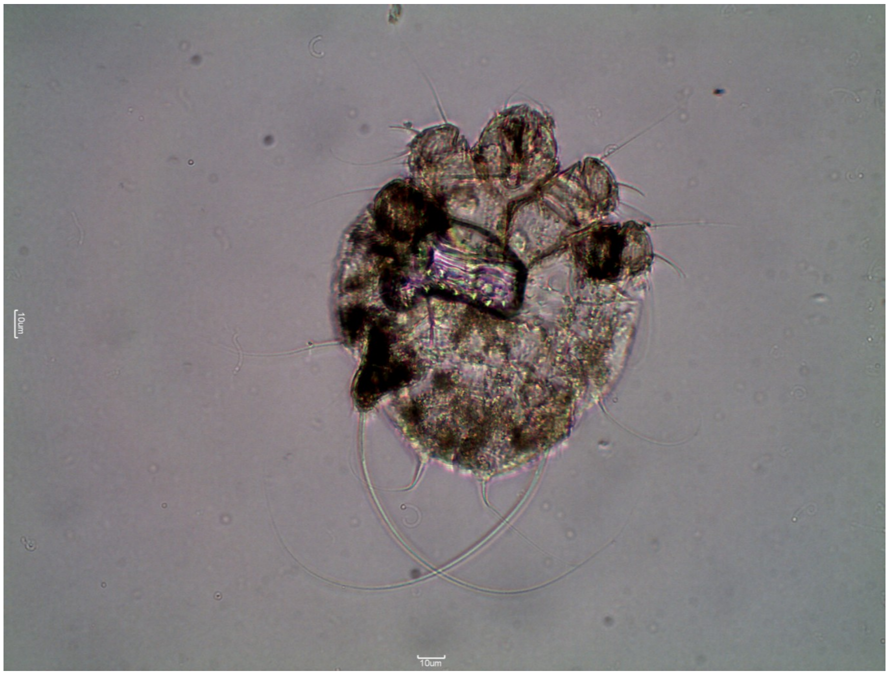

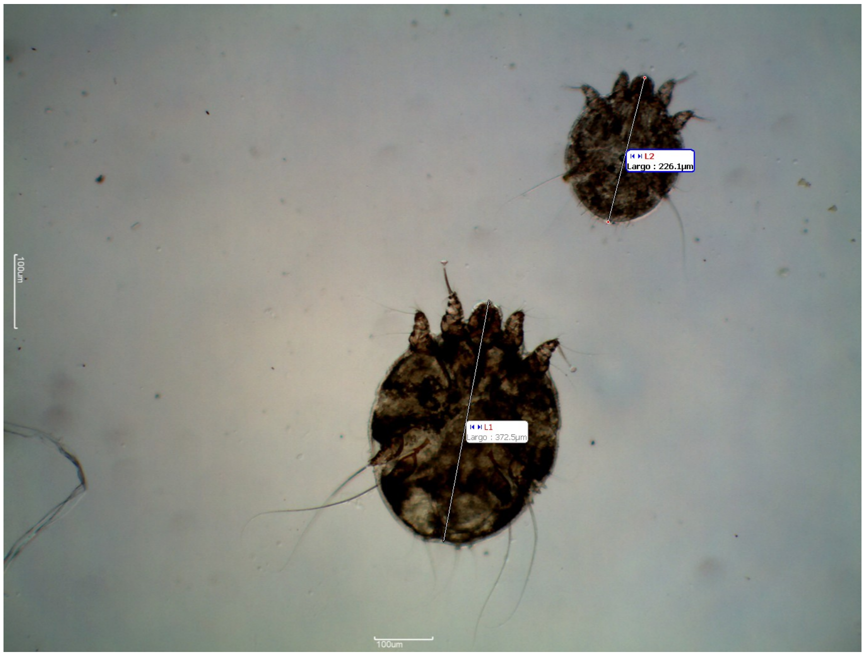
L2
Largo : 226.1µm
L1
Largo : 372.5µm
100um
100um

**Supporting Information 8**

**Provinces containing reports of abnormally alopecic South American camelids provided by park rangers (last three frames as in Figure 1C). The first frame show Provinces with guanacos with confirmed *Sarcoptes scabiei* infestation (green hashed areas) that were reported prior to 2004 (Alvarado, Skewes, and Brevis 2004; Zárate and Valencia 2010).**

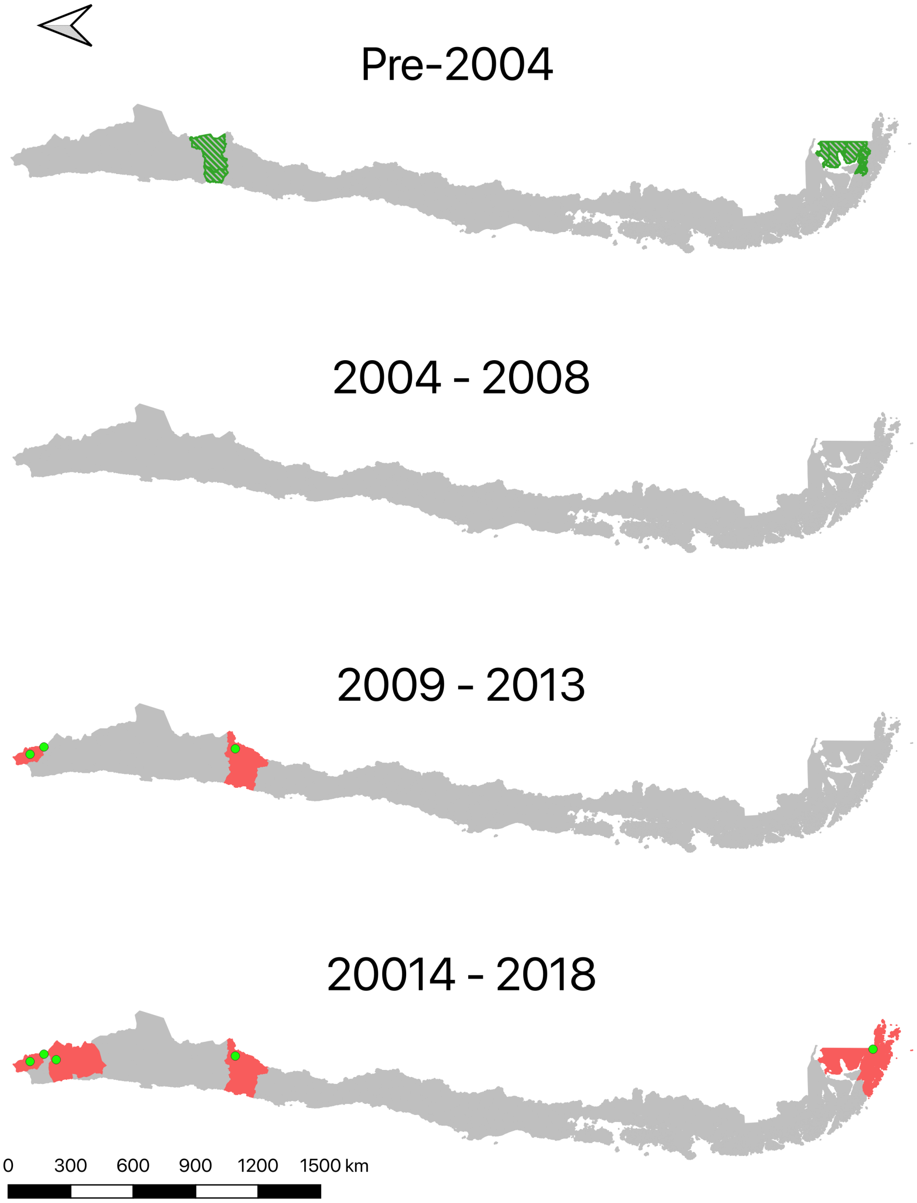
